# Deep learning-based classification of complex intracellular calcium concentration patterns

**DOI:** 10.1101/2025.08.31.673312

**Authors:** Jaesung Choi, Athokpam Langlen Chanu, Shakul Awasthi

## Abstract

Intracellular calcium ion (Ca^2+^) exhibits diverse dynamical behaviors, including complex oscillations such as bursting and chaos. The distinction of diverse dynamical patterns in time series data holds significant biochemical and biophysical implications, as these patterns are intimately related to physiological cellular states associated with health and disease. In this work, we introduce a deep learning framework based on a large kernel convolutional neural network (LKCNN) for the classification of dynamical states of intracellular Ca^2+^ dynamics. We use a chemical Langevin equation to generate synthetic data for training the LKCNN, simulating intracellular Ca^2+^ concentration patterns mimicking real experimental traces. We show that the LKCNN framework reliably classifies diverse intracellular Ca^2+^ dynamical regimes, achieving near 90% accuracy across all dynamical states. To this end, we propose an optimized kernel size for the LKCNN classifier, which captures both short-lived fluctuations and long-range correlations. While steady states, bursting, and simple oscillations are robustly distinguished with near-perfect accuracy, the performance slightly degrades for chaotic and multiple periodicity states under realistic levels of intrinsic fluctuations, reflecting genuine overlap in their temporal signatures. We validate our classifier with experimental Ca^2+^ concentration data, showing strong agreement with manual labeling and confirming generalizability beyond synthetic datasets. These results establish LKCNNs as flexible and scalable tools for systematic classification of cellular dynamics, with broad potential applications to other oscillatory biological processes.

## INTRODUCTION

Patterns are ubiquitous in nature across multiple scales, manifesting in phenomena as diverse as the intricate morphologies of supernovae and galaxies^1,2^, atmospheric turbulence in Earth’s climate system, geophysical flows, and plasma^3^, the collective motion in flocking birds and humans^4,5^, and the rhythmic activity within biological cells^6^. At the cellular scale, calcium ions (Ca^2+^), which serve as crucial intracellular messengers, display rhythmic patterns in their cytosolic concentration when the cell is stimulated by an extracellular agonist^7,8^. These rhythmic patterns are termed as intracellular Ca^2+^ oscillations, and are observed across various cell types, including pancreatic cells^9,10^, hepatocytes^11^, muscle cells^12,13^, and neurons^14^. Intracellular Ca^2+^ oscillations play crucial roles not only in signal transduction inside the cell^15,16^ but also in regulating various physiological processes, including gene expression^17^, cell proliferation^18^, and neuronal differentiation^19^.

Experimental studies observe complex temporal patterns of intracellular Ca^2+^ oscillations. To explain these complex patterns, several mathematical models have been developed, starting from simple minimal models^20^ to complex inositol trisphosphate (InsP_3_) gated models^21–23^. In particular, Houart *et al.*^24^ developed a deterministic model based on the non-linear feedback mechanism of the Ca^2+^-induced Ca^2+^ release (CICR) mechanism^25^, a process prevalent in various cell types, including hepatocytes^26^ and cardiac^27^ cells. In the CICR mechanism, the release of Ca^2+^ from intracellular stores into the cytosol is activated by both InsP_3_ and cytosolic Ca^2+^ itself, forming an autocatalytic feedback loop. This mechanism gives rise to a variety of complex dynamical behaviors, including periodic spikes and bursting in the Ca^2+^ oscillation patterns. Theoretical modeling and experimental observations of these complex Ca^2+^ oscillations are important lines of investigation for a deeper understanding of real Ca^2+^ dynamics in living cells from the perspectives of dynamical systems theory as well as bio-physical and bio-chemical implications in cell biology.

In small biological cells, fluctuations are inherent. Intrinsic fluctuations stemming from random molecular interactions play crucial roles in regulating cellular organizations^28^, cellular decisionmaking, and fate determination^29,30^. Intrinsic fluctuations in intracellular Ca^2+^ oscillations arise due to finite cell size and a small number of reactants^31,32^. One of the present authors^33^ has recently studied the stochastic dynamics of intracellular Ca^2+^ oscillations in Houart’s model^24^ to investigate the behavior of cytosolic Ca^2+^ dynamics driven by intrinsic fluctuations. In this study^33^, a type of entropy known as permutation entropy^34^ (based on ordinal patterns in symbolic dynamics) was proposed to characterize and classify diverse dynamical states of the intracellular Ca^2+^ dynamics. The dynamical states considered include steady-state, simple periodic oscillations, bursting, chaos, multiple periodicity, and quasiperiodic oscillations. At the realistic level of intrinsic fluctuation (hereafter referred to as noise)^35–37^, the permutation entropy’s ability to distinguish states with multiple-periodicity from chaos remains unsatisfactory. In addition, the permutation entropy analysis was based on representative time series from a limited parameter space of the highly nonlinear Houart’s model, and therefore, the generalizability of its conclusions to larger model parameter space (corresponding to broader dynamical regimes) needs to be further investigated. Moreover, to the best of our knowledge, a model-independent classification fit for real experimental data of intracellular Ca^2+^ concentration is not yet known. Hence, it is imperative to develop a robust classifier capable of accurately distinguishing dynamical states of intracellular Ca^2+^ concentration with various noise levels.

The classification of dynamical patterns exhibited by intracellular Ca^2+^ concentration holds practical significance from both bio-chemical and bio-physical perspectives. In Ca^2+^ dynamics, a steady state implies a resting state where the intracellular Ca^2+^ concentration is relatively low ( 0.1–0.2 *µ*M)^38,39^, simple periodic oscillations indicate normal cellular signaling and homeostasis (regulation of accurate control output)^32,40^, while bursting is intimately related to apoptotic cell death^41,42^. Quasiperiodicity has been associated with normal functioning and pathology^43,44^. Period-two or period-three bifurcations are linked with multiple cellular phenotypes in yeast^45^. Further, whether chaos is actually present in biological data and how to accurately characterize them still remain open questions of great interest in non-linear physics of complex systems^46–55^. In general, the characterization of diverse dynamical patterns in simulated/real experimental data of complex biological systems, including Ca^2+^ oscillations, is fundamentally significant as these patterns are closely linked with physiological states related to health and disease^56,57^.

In this work, we propose using machine learning (ML) to classify the dynamical states of intracellular Ca^2+^ concentration. ML has been widely applied to the analysis of biological processes, including diffusion dynamics in single-particle tracking trajectories^58–60^ and gene expression classification from microarray data^61^. The traditional ML approaches, however, often face significant challenges with biological data, which are often noisy, heterogeneous^58^, and have limited data points. Further, traditional ML approaches rely on extensive manual feature engineering, which proves to be difficult for such biological data^62^. These difficulties, therefore, hinder ML classification performance. Consistent with this, the experimental data of intracellular Ca^2+^ concentration also face these challenges: experimental traces of Ca^2+^ concentration are not only noisy but also exhibit complex temporal patterns such as bursting, making manual feature extraction highly challenging for accurate classification.

Recent advances in deep learning offer promising solutions to the challenges mentioned above, automatically learning complex temporal patterns directly from raw data, eliminating manual feature engineering requirements^63,64^. A fundamental challenge in applying deep learning to Ca^2+^ dynamics is in obtaining accurate labels for training. This necessitates the use of well-validated synthetic datasets generated from mathematical models to train robust classifiers that can subsequently be applied to experimental data. For this transfer to succeed, the model must effectively generalize dynamical features across different data domains. To achieve this, we employ the large kernel convolutional neural network (LKCNN), which demonstrates superior dynamical feature generalization capabilities for characterization of time series generated by complex systems^65–67^. While conventional deep learning architectures such as residual networks (ResNets) often struggle with the temporal complexity inherent in dynamical systems, LKCNN has demonstrated robust generalization capabilities specifically for complex system characterization, achieving 89.8% accuracy^65^. Unlike standard CNNs with small kernel sizes (∼3 − 5)^68^, LKCNN relies on large-scale observational windows, enabling it to capture long-range correlations and complex temporal patterns of various dynamical behaviors. Recent analysis suggests that LKCNN appears well-suited for learning qualitative dynamical features beyond simple periodic patterns, which may explain its superior ability to distinguish between chaotic and regular behaviors across different dynamical systems^66^.

In this study, we present a deep learning framework specifically designed for intracellular Ca^2+^ dynamics classification. We extend beyond binary classification approaches used in previous LKCNN studies and distinguish multiple dynamical states: steady states, simple periodic and multiple periodicity oscillations, bursting, chaotic (aperiodic), and quasiperiodic patterns. Although advanced variants such as the Siamese large kernel convolutional neural network (SLKCNN) have been proposed^69^, we adopt the original LKCNN architecture as a prototype to investigate the intracellular Ca^2+^ dynamics. Our approach leverages synthetic data from the well-known Houart’s nonlinear model of intracellular Ca^2+^ oscillation in order to train LKCNN classifiers that demonstrate robust generalization to real experimental Ca^2+^ concentration time series data. This methodology enables automated analysis of complex temporal patterns crucial for understanding cellular physiological states and disease mechanisms, eliminating the need for manual feature engineering and expert-dependent pattern recognition.

## RESULTS

### Intracellular Ca**^2+^** concentration exhibits complex dynamical patterns

The training of a neural network typically requires large and high-quality datasets with true labels. Since obtaining accurate labels directly from noisy experimental time series data is challenging, we employ a simulation-based approach for training our LKCNN model. One may use a variety of theoretical models that can reliably reproduce the experimentally observed complex temporal patterns of intracellular Ca^2+^ dynamics. In this work, we choose the nonlinear model presented by Houart *et al.*^24^ since it can mimic various dynamical states, hence making it suitable for a comprehensive training and classification of intracellular Ca^2+^ dynamics. The nonlinear model centers around the interplay between Ca^2+^-induced Ca^2+^-release (CICR) and Ca^2+^-activated 1,4,5-trisphosphate (InsP_3_) degradation mechanisms, where the coupling of a negative feedback loop (degradation) with a positive CICR cycle gives rise to complex intracellular Ca^2+^ oscillations^26^. A schematic representation of this mechanism is shown in panel (a) of Fig. 1, and we detail the governing theoretical nonlinear model in the Methods Section (Eq. (3)).

**Figure 1:**
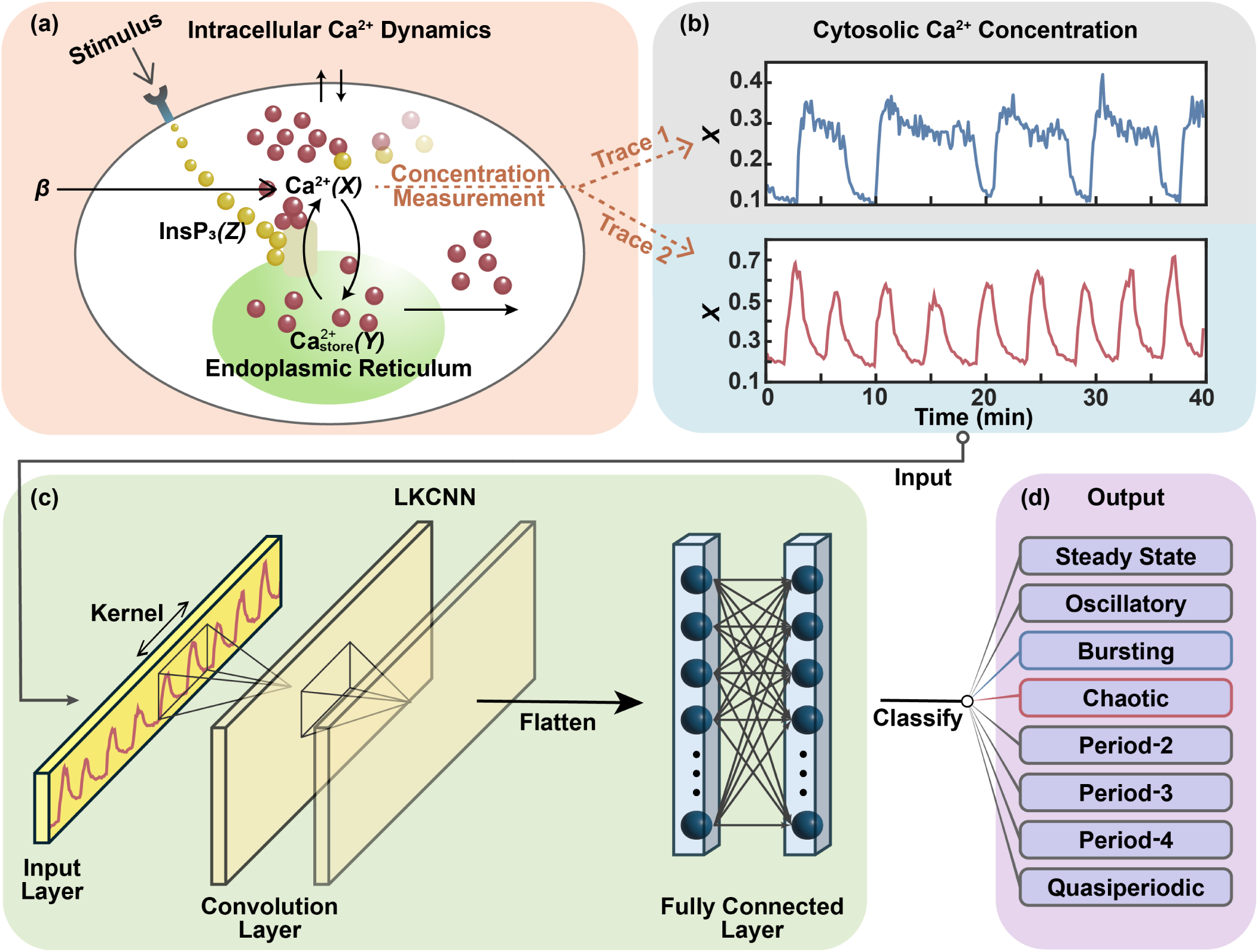
Workflow diagram: Our approach processes diverse dynamical patterns of intracellular calcium (Ca^2+^) dynamics in simulated trajectories as well as experimental traces. Simulated data are generated by the interplay between the mechanisms of Ca^2+^-induced Ca^2+^-release (CICR) and Ca^2+^-activated 1,4,5-trisphosphate (InsP_3_) degradation inside a biological cell as illustrated in (a). See Methods section for detailed descriptions of the intracellular Ca^2+^ oscillation model. Experimental traces of Ca^2+^ concentration are obtained from different cells (see the Main text). First, we generate simulated cytosolic Ca^2+^ concentration trajectories from the intracellular Ca^2+^ oscillation model or obtain experimental traces from publicly available datasets (two representative traces shown in (b)). We then feed the simulated trajectories or experimental traces to the large kernel convolutional neural network (LKCNN) model (as shown in (c)), which classifies the diverse dynamical patterns of cytosolic Ca^2+^ concentration into correct labels (d).

It is well-known that deterministic models, such as the one given in Eq. (3), deal with continuous concentrations by assuming an infinitely large number of molecules. Thus, these models prove insufficient for systems with low molecule counts, such as those at an intracellular scale. Ideally, the concentration would be replaced by a discrete number of molecules. However, this approach proves to be difficult to handle both analytically and numerically. To alleviate this problem, we adopt an intermediate strategy that allows fluctuations in concentration by explicitly adding stochastic terms; the resultant equation is known as the chemical Langevin equation (CLE)^70,71^. Following this approach, we derive the CLE pertaining to the intracellular Ca^2+^ oscillation model (see Methods Section for a detailed derivation) as follows:

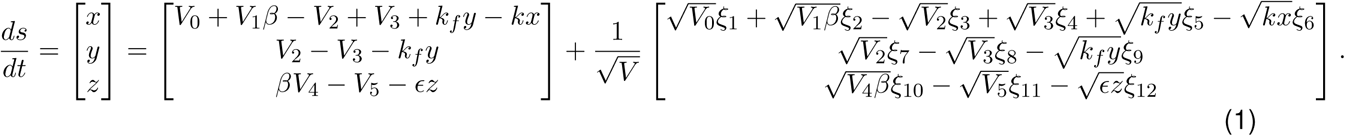

where *ξ_j_* (*j* = 1, 2*,…,* 12) denotes mutually independent Gaussian white noise processes characterized by ⟨*ξ_j_*(*t*)⟩ = 0 and ⟨*ξ_j_*(*t*) *ξ_j_′* (*t^′^*)⟩ = *δ_jj_′ δ*(*t* − *t^′^*). While the first term of Eq. (1) is simply the deterministic nonlinear model (Eq. (3)), the second term represents the stochastic component of the intracellular Ca^2+^ dynamics. The prefactor 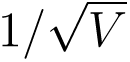 explicitly accounts for the influence of system size (*V*) or intrinsic fluctuations on the evolution of Ca^2+^ concentration. Specifically, the parameter *V* determines the amplitude of intrinsic fluctuations: as *V* increases, the relative contributi<u>o</u>n of stochastic effects diminishes. In the thermodynamic limit, where *V* → ∞, the term 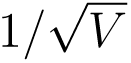 tends to zero, causing intrinsic fluctuations to vanish altogether. These intrinsic fluctuations manifest as inherent noise in the underlying dynamics, making classification difficult. Numerically solving the CLE (1) using the Euler-Maruyama method for different values of the rate constants and other model parameters (detailed in the Methods Section), concentrations *x*, *y*, and *z* in Eq. (1) exhibit various kinds of dynamics. For visual reference, representative trajectories are presented in Fig. 2, which illustrates characteristic temporal patterns in cytosolic Ca^2+^ concentration *x*(*t*) (expressed in *µ*M) at three levels of noise *V* =, 10^5^ and 10^3^. The panels illustrate: (a) steady state, (b) simple periodic oscillations (oscillatory), (c) bursting, (d) chaotic, (e) period-2 (P-2), (f) period-3 (P-3), (g) period-4 (P-4), and (h) quasiperiodic oscillations. Time *t* is measured in minutes (min).

**Figure 2:**
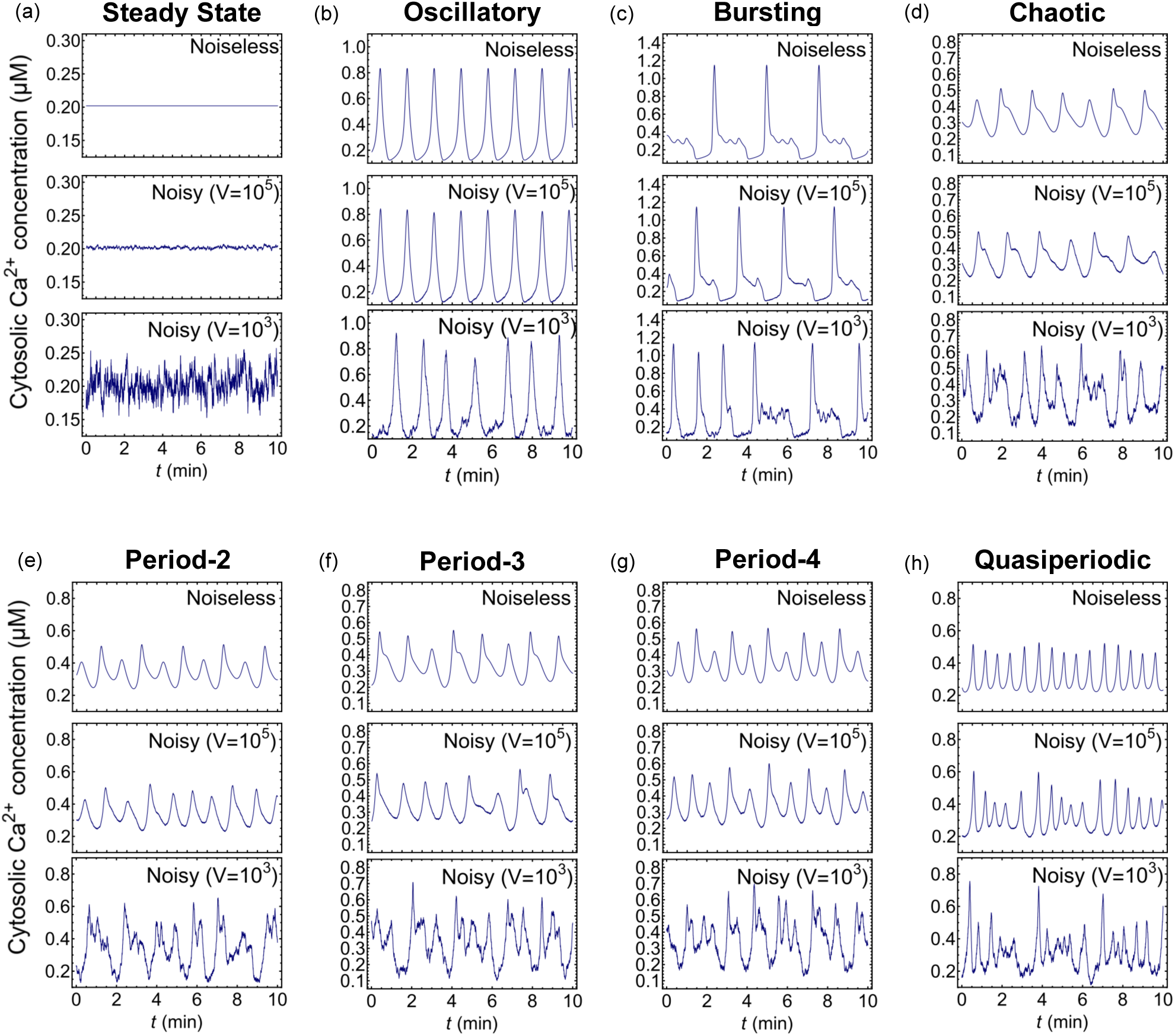
**Various dynamical states of cytosolic Ca**^2+^ **concentration:** (a) Steady state, (b) oscillating state, (c) bursting, (d) chaotic state, sequences of (e) period-2, (f) period-3, (g) period-4, and (h) quasiperiodic oscillations. For each panel, the three rows correspond to deterministic or noiseless (*top*), noisy with *V* = 10^5^ (*middle*), and *V* = 10^3^ (*bottom*).

We use the dynamical states of cytosolic Ca^2+^ concentration *x*(*t*) as inputs for our LKCNN classification (as illustrated in panels (b) and (c) of Fig. 1). Physiological bounds on cytosolic Ca^2+^ concentration, typically from below 0.1 *µ*M up to 1 *µ*M with oscillation periods ranging from a few seconds to ~30 min^24,72^, limit the parameter space of interest. The model in Eq. (1) however contains 19 tunable parameters whose variation can generate a wide range of dynamical states. For training and testing our neural network, we therefore consider three parameter regions that together largely cover the observed amplitude and frequency range of Ca^2+^ oscillations. These regions have been numerically analyzed previously in the original study^24^, and we provide the exact values of the parameters used in the Methods section. However, in our study, we further subdivide the parametric region close to chaos into multiple-periodicity states, namely period-2 (P-2), period-3 (P-3), and period-4 (P-4) oscillations to better highlight the advantages of the deep-learning approach.

### An optimal LKCNN architecture is obtained for robust classification of intracellular Ca**^2+^** dynamics

We employ the LKCNN architecture shown in panel (c) of Fig. 1 to classify the various dynamical states of intracellular Ca^2+^ dynamics at varying levels of noise (quantified by *V*), leveraging its demonstrated effectiveness in dynamical systems analysis^65,66^. However, prior to training and classifying the dynamical states, we have to properly initialize the neural network. The initialization is set through the seed, which determines the weights of the LKCNN before training. Our analyses indicate that the choice of seed has only a minimal impact on the overall classification performance, highlighting the robustness of the LKCNN classifier. The other parameter is the kernel size *k*, which is defined as the network’s temporal receptive field. It is the span of consecutive data points of a time series that it analyzes simultaneously to detect patterns. This value influences how much local context the neural network considers when extracting features from the input data. Previous studies have demonstrated the effectiveness of LKCNN by using a fixed kernel size of 100 for the binary classification of chaotic versus regular dynamics in noise-free systems^65,66^, showing that large kernels enable the direct capture of extended temporal dependencies and long-range correlations that characterize complex dynamical behaviors. For instance, long-range patterns like P-3, P-4, and bursting necessitate a large temporal receptive field for effective detection. However, a trade-off exists: an excessively large kernel, while beneficial for capturing these long-term dependencies, can obscure the fine-grained details of shorter-term patterns by blurring their local features. Since the amplitude, frequency, and noise levels of the Ca^2+^ concentration dynamics all lie within well-defined, experimentally-determined ranges, the kernel size used for classifying the dynamical patterns is also expected to perform optimally only within a specific range. Therefore, identifying an optimal kernel size that is large enough for long-range correlations yet fine enough to preserve local details is crucial for accurately distinguishing the full spectrum of Ca^2+^ dynamical behaviors. Operationally, the LKCNN architecture proceeds in such a way that the large kernel slides across the input Ca^2+^ concentration time series to extract temporal features through successive convolutional layers and the extracted features are then flattened and processed through fully connected layers to produce the final classification output (as illustrated in Fig. 1(d)). The detailed LKCNN architecture and training procedure are described in the Methods section.

We now analyze the performance of the LKCNN classifier across a range of kernel sizes *k* under two distinct scenarios of cytosolic Ca^2+^ concentration: (i) an ideal, noiseless condition, and (ii) a more realistic, noisy condition. This allows us to investigate the impact of noise on the classifier’s performance as well as to identify a range of kernel sizes that demonstrate robustness in noisy conditions. In the first setup, which we refer to as the “Noiseless” scenario, we evaluate the performance of the classifier on the noise-free (*V* = ∞, deterministic dynamics) test data. The second setup referred to as “Noisy” scenario assesses our classifier’s robustness against test data with a wide range of *V* values necessarily including the realistic noise levels (*V* ɛ 10^6^, 10^5^)^33,35–37^ of intracellular Ca^2+^ dynamics. For both setups, we train the LKCNN on a dataset spanning noise levels *Vɛ*{∞, 10^8^, 10^7^, 10^6^, 10^5^}. For each value of *V*, the training dataset is created using at least 1,000 trajectories from each of the eight dynamical states of Ca^2+^ dynamics. Similarly, another dataset with 12,000 trajectories was generated to test the performance of the classifier. Since our synthetic datasets contain time trajectories of various dynamical states (hereafter also referred to as labels or class) of cytosolic Ca^2+^ concentration, we quantify our classifier’s performance as its ability to solve a multi-class classification problem, hence termed classification accuracy. That is, the machine learning task is to predict and correctly assign one of the eight distinct dynamical pattern labels to each test data sample. We define the classification accuracy as the fraction of samples for which the LKCNN classifier’s predicted label exactly matches the true label as:

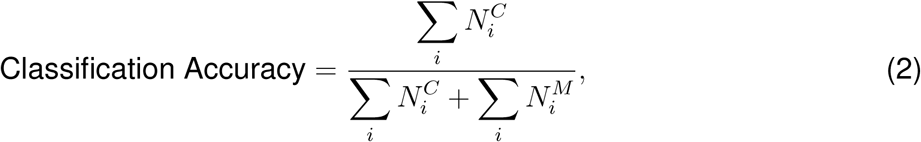

where *N_i_^C^*and *N_i_^M^*represent respectively the number of correctly classified and misclassified samples *i* (steady state, bursting, oscillatory, P-2, P-3, P-4, chaotic, and quasiperiodic oscillations).

To ensure reproducibility and determine the optimal architecture, we train and subsequently test the accuracy of the LKCNN classifier with various seeds and kernel sizes. Fig. 3 presents the classification accuracy for the noiseless (blue) and noisy (orange) test datasets against the kernel size *k* of the LKCNN. We take 20 different random seeds and *k* = 4, 5, 6*,…,* 100 (indicated by circle and square markers). The solid lines indicate the mean accuracy averaged over the multiple seeds. The shaded envelopes denote one standard deviation (±*σ*), indicating the stability of our LKCNN classifier’s performance. We observe a narrower band of blue shaded envelope under noiseless conditions as compared to the noisy condition at *k ≥*30, implying a more stable performance for the noise-free or deterministic dynamics of cytosolic Ca^2+^ concentration across different random seeds. We further observe that the presence of noise significantly lowers the multi-class classification performance of the classifier, indicated by the lower accuracy values of the noisy data condition compared to the noiseless case. Notably, we observe that the two cases show distinct relationships between the LKCNN kernel size and classification accuracy. In the noiseless scenario, the accuracy (blue curve) rapidly improves with an increase in kernel size, followed by a saturation around *k* 30. On the other hand, for the noisy data condition, a more realistic scenario, the accuracy curve (orange) shows a clear peak for kernel sizes ~ 25– 30, indicating an optimal classification performance of the LKCNN within this range. Beyond this range, the performance of the classifier steadily declines, which indicates that while larger kernels are beneficial for the noiseless data condition, moderately-sized LKCNN kernels are optimal for datasets with realistic noise levels. We thus obtain an optimal LKCNN architecture for robust classification of the intracellular Ca^2+^ dynamics from synthetic numerically-simulated trajectories that can be used for analysis of real experimental Ca^2+^ concentration traces. For all subsequent analyses, we use a kernel size of *k* = 28 because it yields the best performance in both noisy and noise-free settings.

**Figure 3:**
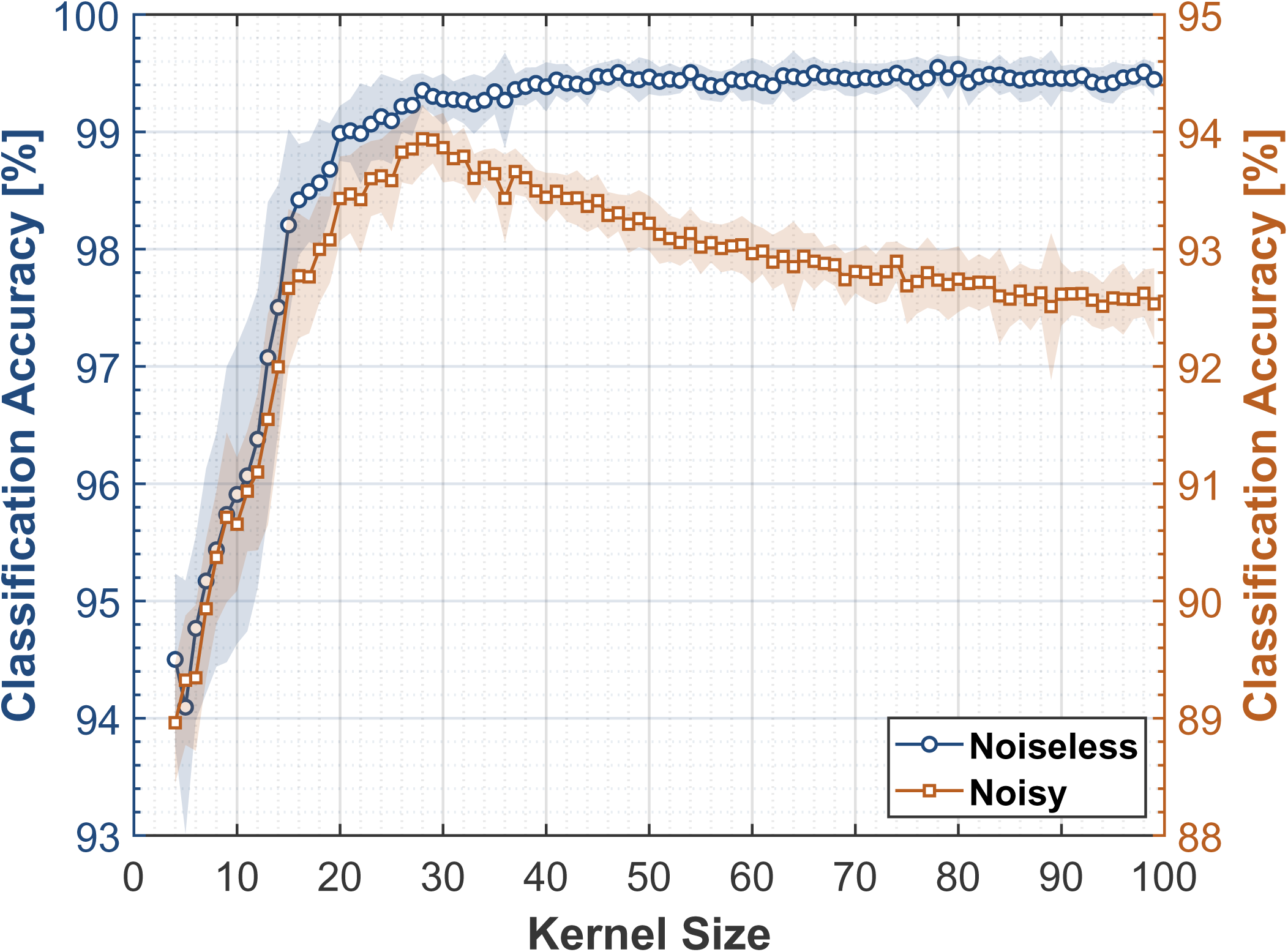
Accuracy of LKCNNs over a range of kernel size for noiseless (blue curve) and noisy (orange curve) trajectories of cytosolic Ca^2+^ dynamics. The shaded region denotes one standard deviation (±*σ*).

### LKCNN shows high classification accuracy in the data of noiseless Ca**^2+^** states

Having obtained the optimal LKCNN classifier, we now analyze its performance at the level of individual dynamical states. Rather than relying solely on aggregate accuracy, we report per-state accuracy, precision, recall, and F1 scores to reveal any heterogeneity across the eight states. We further examine the confusion matrix to identify systematic misclassifications and shared error modes between closely related patterns. To probe robustness, we stratify these metrics by noise level *V*, assessing which states are most sensitive to increased noise.

We focus on the noiseless case in this section. We continue with the optimized LKCNN classifier trained on the noise levels of *Vɛ* {∞,10^8^, 10^7^, 10^6^, 10^5^} with *k* = 28, and subsequently evaluate its performance on a distinct test set of 12, 000 noiseless trajectories. Table 2 reports class-wise performance, indicating a uniformly high accuracy of 99.4%. Precision, which is defined as the proportion of predicted class instances that are correct, recall, which is the proportion of true class instances recovered, and F1 score, which is the harmonic mean of precision and recall, are near unity for most states. Steady state, bursting, oscillatory, and quasiperiodic are effectively identified. The multiple-periodicity states P-2, P-3 and P-4 are discerned with high precision but slightly lower recall (∼0.969–0.996), yielding F1 scores of 0.985 (P-2), 0.982 (P-3), and 0.981 (P-4). Performance is comparatively weaker for chaotic, suggesting a tendency toward false positives in this class. Overall, the LKCNN works effectively as a classifier for all states, with minor errors concentrated in chaotic and, to a lesser extent, P-3 and P-4 states. The numerical values of these metrics for the eight dynamical states are given in Tab. 2.

We continue our analysis with the confusion matrix shown in Fig. 4 in which the rows correspond to the true labels of the simulated cytosolic Ca^2+^ concentration trajectory data, while the columns represent the labels predicted by our LKCNN classifier. The diagonal entries (blue colored) show the percentage of each class that has been correctly identified (per-class accuracy), whereas off-diagonal values indicate misclassifications. Our result shows that the optimized LKCNN achieves a high level of classification performance on the noiseless test data, achieving an overall average accuracy of 99.5%. The diagonal values, many of which are nearly 100.0%, thus indicate near-perfect classification for most dynamical states of cytosolic Ca^2+^ concentration, including steady state, bursting, oscillatory, P-3, P-4, and quasiperiodic oscillations. We see that a small fraction of P-2 samples are confused with P-4 (2.0%), and a small fraction of chaotic trajectory samples are misidentified, primarily as P-3 (2.2%). These misclassifications suggest that while our LKCNN classifier is highly effective, its most significant challenge lies in distinguishing between the fine-grained features that separate chaotic behavior from certain multiple-periodicity states. Nonetheless, the high classification accuracy shown on the diagonal of the confusion matrix demonstrates the excellent distinguishing power and reliability of our LKCNN classifier in noise-free datasets.

**Figure 4:**
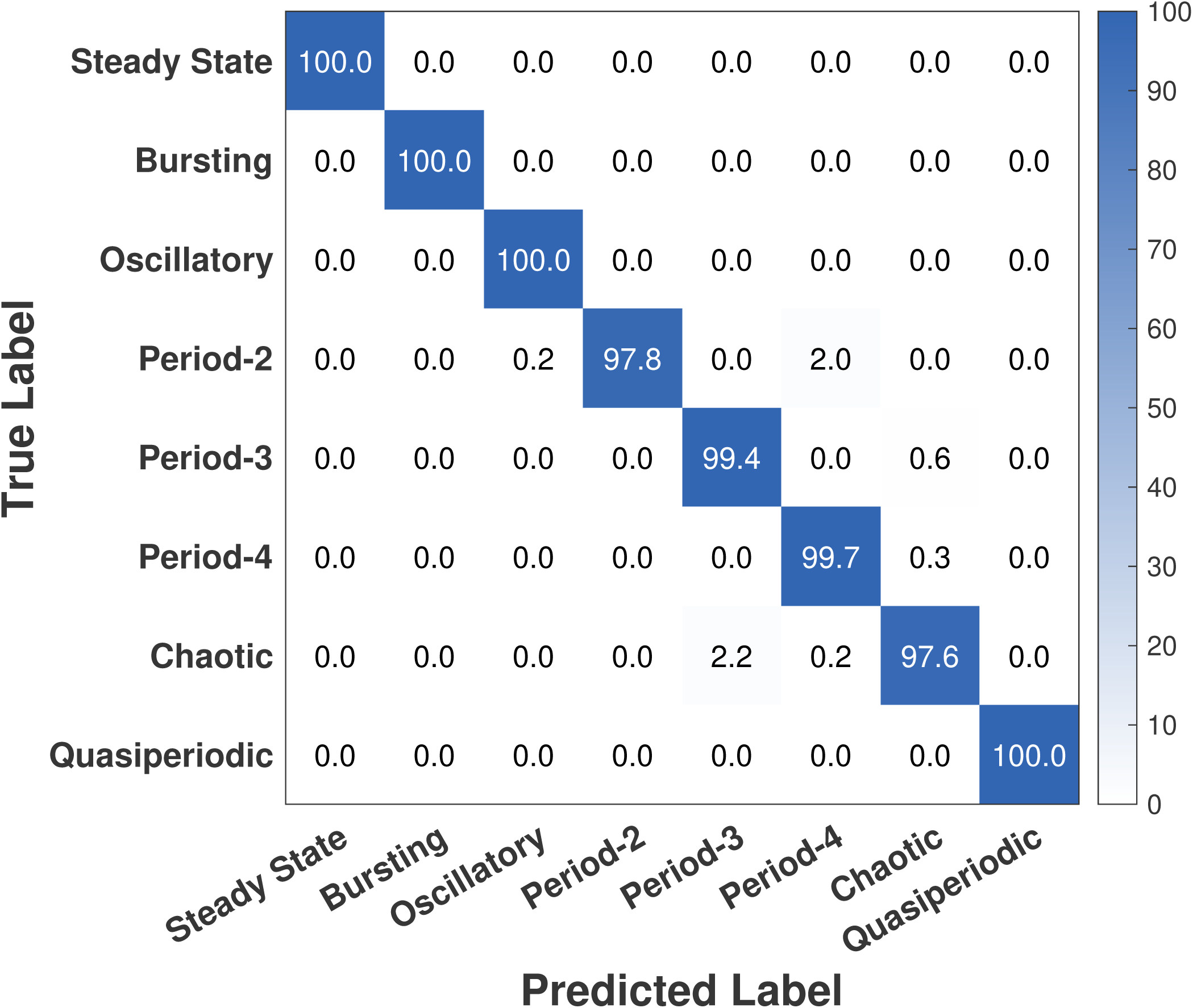
Confusion matrix showing the result of multi-class classification performance of our LKCNN model for the noiseless case. Diagonal entries denote the percentage of correctly predicted dynamical states of cytosolic Ca^2+^. Labels indicate steady state, bursting, oscillatory, period-2, period-3, period-4, chaotic state, and quasiperiodic oscillation.

### Noise degrades the performance of LKCNN for classifying chaotic and multi-periodicity Ca**^2+^** dynamical states

We now extend our analysis to examine the performance of the optimized LKCNN classifier at the level of individual dynamical states in the presence of noise. For a more comprehensive analysis of the noise effect, we extend both the training and test datasets to cover increasingly noisy conditions, corresponding to smaller *V* values. While the kernel size *k* is fixed at 28 as stated earlier, the present configuration is fundamentally different because training is performed across a wider range of noise levels, *Vɛ*{∞,10^8^, 10^7^, 10^6^, 10^5^, 10^4^, 10^3^}. A broader and more extreme spectrum of noisy conditions is considered to improve the robustness of the LKCNN classifier and better capture the dynamical behavior of Ca^2+^ states under challenging test scenarios. This setup also allows us to investigate performance over more distinct noise levels than before.

Figure 5 shows how the classification accuracy of the LKCNN classifier varies with the cumulative noise level in the test dataset. Here, “cumulative” means that the noise level *V*, plotted on the *x*-axis, represents the maximum noise level included in the test set. For example, *V* = 10^6^ indicates that the test set contains all samples from *Vɛ*{∞,10^8^, 10^7^, 10^6^}. Similarly, the accuracy value at *V* = 10^3^ is computed using the entire test dataset from *V* = through *V* = 10^3^. The cumulative test set mimics a realistic condition by progressively broadening the distribution of noise in the evaluation data, as in experimental scenarios, a deployed classifier would not see only one fixed noise level, and would face a mix of conditions. The green circles in Fig. 5 denote the mean accuracy over 20 runs with different random seeds for each cumulative noise level, while the shaded green region indicates the corresponding standard deviation (±*σ*). The highest accuracy (≈98.5%) occurs when testing on the noiseless dataset (*V* =∞). As noise is progressively added, mean accuracy steadily declines, remaining relatively high (≳ 98.0%) for *V≥*10^6^, but dropping more noticeably when *V≤*10^5^, reaching ≈91.0% at *V* = 10^3^. These results show that the quality of the input data inherently constrains the LKCNN’s classification performance. Note that for *V* =∞, the accuracy of 98.5% is slightly below the *>* 99.0% achieved in Fig. 3, as the training set for the latter was limited to *Vɛ*{∞,10^8^, 10^7^, 10^6^, 10^5^}. This suggests that including extremely noisy data (*V* = 10^3^) in training slightly degrades performance even when testing on noiseless data. Indeed, when training is restricted to noise levels up to *V* = 10^4^ (excluding *V* = 10^3^), performance on the noiseless test set almost matches the results in Fig. 3.

**Figure 5:**
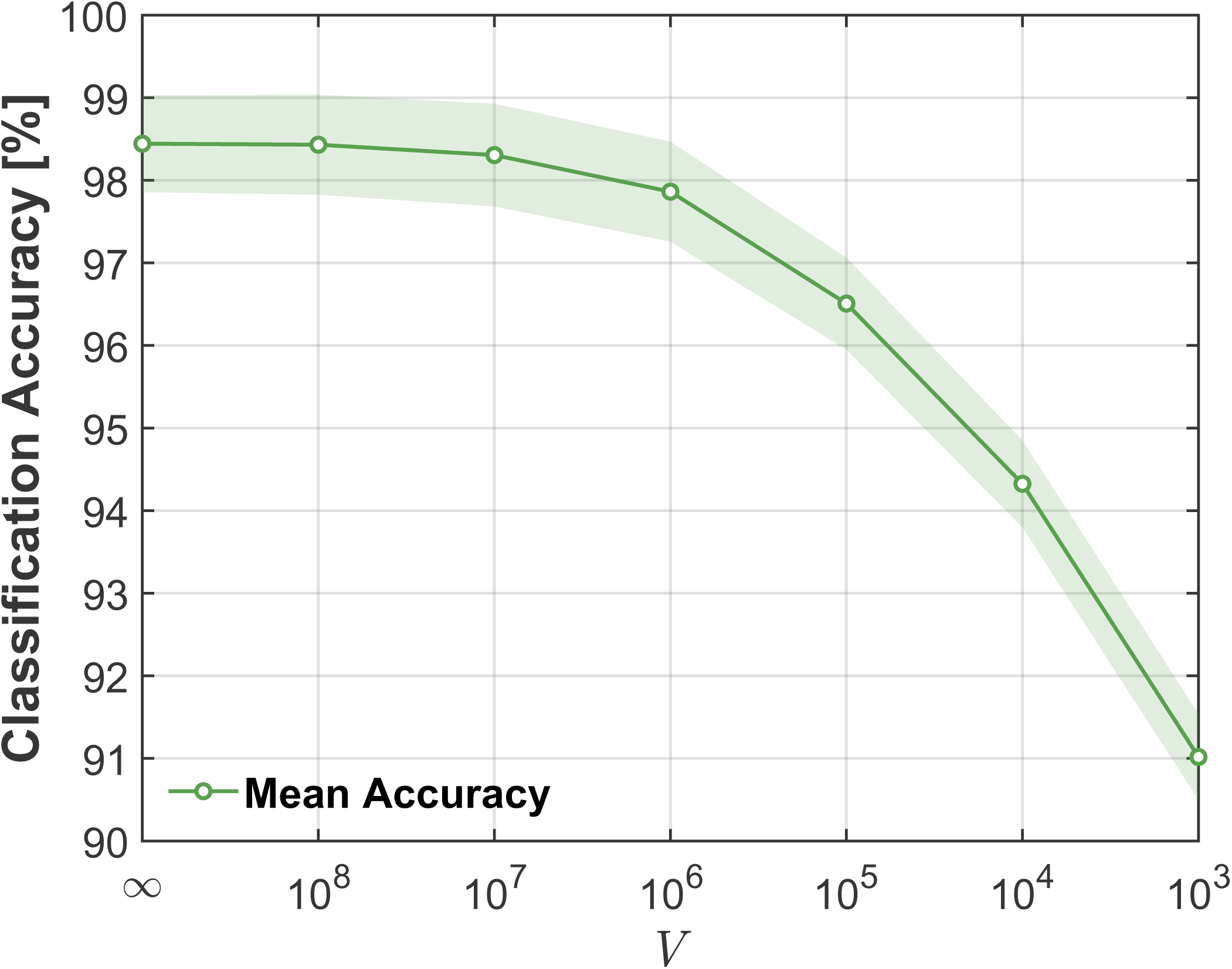
Accuracy versus noise (∝ 1/*V*) in the cytosolic Ca^2+^ dynamics data.

Restricting our subsequent analysis to realistic noise levels in Ca^2+^ dynamics (*Vɛ*{10^6^, 10^5^}^33^) at which the classification accuracy remains above 96%, we ensure the evaluation reflects physiologically relevant conditions. We use the same optimal LKCNN with kernel size *k* = 28 as before. We construct the training dataset by combining 12, 000 samples from each of the *V* levels ({∞, 10^8^, 10^7^, 10^6^, 10^5^}), resulting in a total of 60, 000 training samples. The test set is then constructed with the exact same distribution as the training data, also containing 12, 000 samples from each of the five *V* levels for a total of 60, 000 test samples. Table 3 summarizes the classwise performance on the noisy set, showing high but non-uniform accuracy (overall = 94.00%). Bursting is essentially perfectly identified, and steady state, oscillatory, and quasiperiodic also perform good (F1 scores ≳ 0.980). Performance is lower for the multi-periodicity states P-2, P-3, and P-4 (F1 scores ≈ 0.840–0.860), with P-2 mainly limited by precision, and both P-3, P-4 mainly limited by recall. The weakest class is chaotic (F1 ≈ 0.781), indicating substantial confusion with other labels. Overall, LKCNN remains robust under noise for most behaviors, with errors concentrated in chaotic and, to a lesser extent, the multi-periodicity states. Moreover, on comparing with the metrics for the noiseless data, we note a clear reduction in the overall accuracy of approximately 5.41%. An examination of the F1 scores across individual classes reveals that the most substantial performance degradation occurs in the chaotic class. Other classes exhibiting notable decreases include P-2, P-3, and P-4. These findings suggest that such classes are particularly sensitive to the introduction of noise, possibly due to the perturbation of key discriminative features within their representations. In contrast, several classes demonstrate substantial robustness to noise. The steady state, for example, experiences only a marginal reduction in F1 score from 1.000 to 0.996. Similarly, the bursting and quasiperiodic classes maintain relatively stable performance under noisy conditions, indicating that their underlying feature patterns remain discernible despite the fluctuations.

Fig. 6 presents the confusion matrix for test datasets containing realistic noise levels of intracellular Ca^2+^ dynamics, where we find an overall average accuracy to be 97.42%. While this value is still a high accuracy, it represents a noticeable drop from the previous 99.54% achieved in the noiseless case (Fig. 4). A detailed breakdown of the confusion matrix of Fig. 6 shows how this performance degradation is distributed across the different dynamical classes. We see that the diagonal entries (orange colored) of some of the classes are lower than those observed for the noiseless case (Fig. 4), indicating an increase in misclassifications for these classes. We state some key findings from Fig. 6 as follows. The confusion among different multi-periodicity states has worsened from before. For example, 6.8% of P-2 samples are incorrectly labeled as P-4. The confusion between chaos and periodic states (P-2, P-3, and P-4) has noticeably increased. A significant 8.6% of true chaotic samples are now misclassified as P-3. Conversely, 5.0% of P-3 samples and 3.0% of P-4 samples are misidentified as chaotic. These results demonstrate that the high accuracy observed in the case of noiseless datasets is significantly affected when noise is present, particularly between chaotic and multiple-periodicity states.

**Figure 6:**
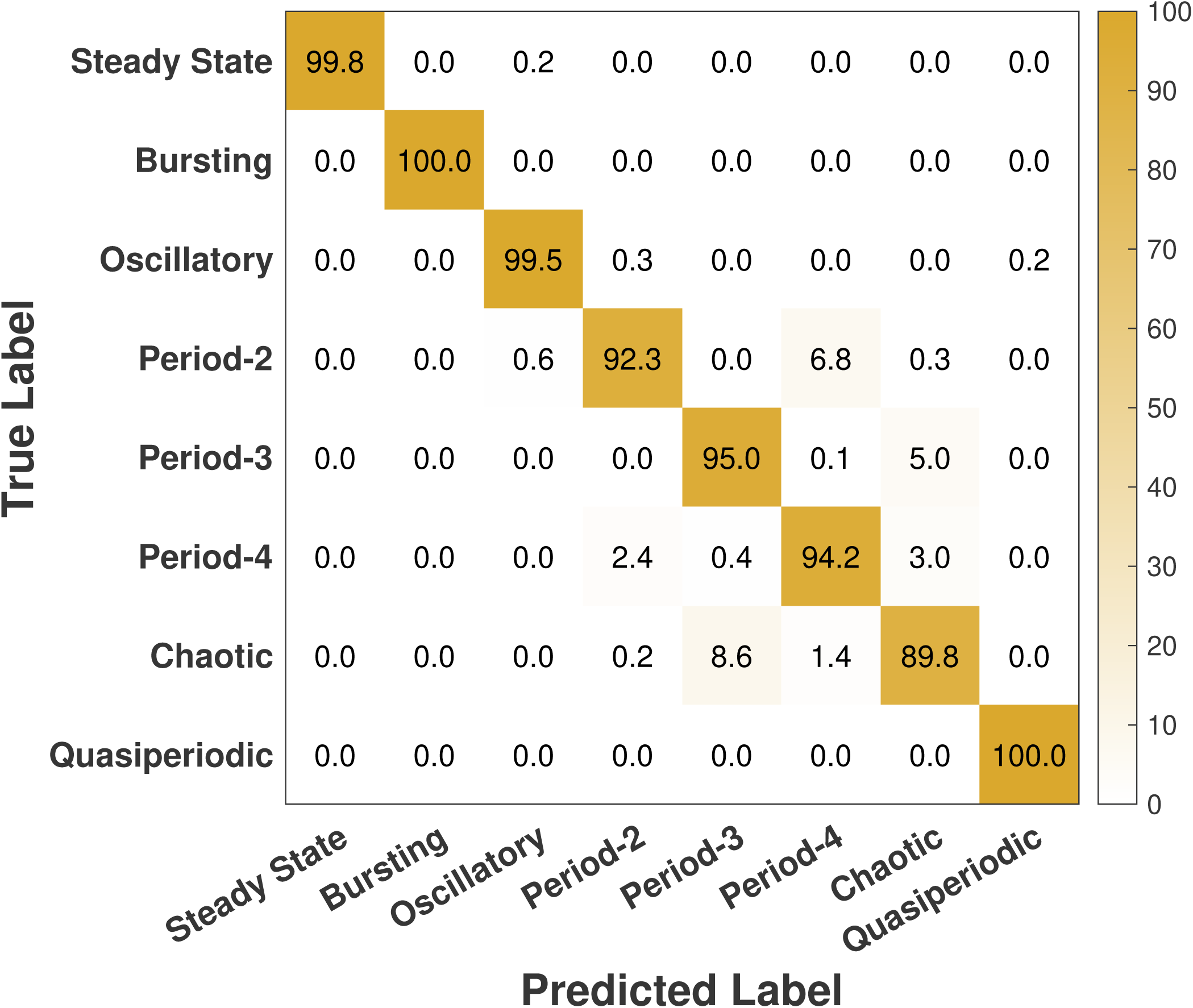
Confusion matrix showing the result of multi-class classification performance of our LKCNN model for the case of realistic noise levels. Diagonal entries denote the percentage of correctly predicted dynamical states of cytosolic Ca^2+^. Labels indicate steady state, bursting, oscillatory, period-2, period-3, period-4, chaotic state, and quasiperiodic oscillation.

### The optimized LKCNN demonstrates generalization capability to experimental data

Having trained and validated the LKCNN classifier exclusively on synthetic datasets, we now turn to its deployment for practical use. In order to bridge the gap between model development and real-world applicability, we use the optimized and tested classifier to process experimental measurements as input and assign them to the appropriate classes. While the network has been exposed only to controlled synthetic data during training, deploying it on experimental data allows us to evaluate its generalization capacity and robustness under realistic conditions.

To evaluate the generalizability of our optimized LKCNN classifier, we now construct a dataset of Ca^2+^ concentration experimental data from two sources: (a) pancreatic islets of mouse *β*-cells^73,74^ displaying diverse patterns in Ca^2+^ concentration recorded in eight genetically distinct mouse strains, and (b) WT-HEK293, STIM-KO, and ORAI TKO cells^75,76^ exhibiting a diversity of Ca^2+^ traces upon different agonist stimulation conditions, like varying concentrations of carbachol (CCh). In the absence of any physiologically motivated ground-truth classification, we rely on manual labeling of experimental traces wherever possible. From the available dataset, we conduct a careful visual screening of the time series to identify clearly distinguishable dynamical regimes. While the dataset comprises a large number of time series, only those with unambiguous dynamical features are deemed suitable for classification. The characteristic features of bursting dynamics include a plateau fraction (time spent in the active oscillation phase) followed by a silent duration phase in the temporal pattern. Steady states are characterized by relatively low (∼0.1 − 0.2*µ*M) concentration. However, it proves difficult to visually identify all other states due to the noise levels as well as the length of data available in the experimental data sets. Following this criterion, we identify 46 time series in the bursting class and 39 time series in the steady state class, and 41 are manually classified as “others” and include visually indistinguishable classes like chaos, oscillatory, quasi-periodic and the multiple-periodicity states.Therefore, the curation process yields a total of 126 in three classes, namely (i) steady state, (ii) bursting, and (iii) others. Since the dataset contains only Fura-2 ratio measurements of intracellular Ca^2+^ concentration, we have employed the Grynkiewicz equation^77^ with estimated calibration parameters to convert the ratio data into approximate Ca^2+^ concentrations. Moreover, we have used linear interpolation to make the experimental traces of exactly 1000 data points, making it suitable for the optimized LKCNN. Using this procedure, we create a benchmarking dataset for validating the performance of our LKCNN classifier on experimental Ca^2+^ concentration dynamics.

When applied to the experimental dataset, the LKCNN has successfully produced stable classifications across all inputs. Representative examples of the experimental trajectories, together with their correctly classified labels, are shown in panels (a), (b), and (c) of Figure 7. Out of the entire set of 126 time series, only one bursting trajectory and three trajectories marked as others were misclassified, resulting in an overall accuracy of 96.8%. These four misclassified trajectories are shown in panel (d) of Figure 7 with their LKCNN-predicted labels indicated by the text in red color. While the trajectory in the upper panel is manually labeled as bursting, LKCNN predicts it as a chaotic state. The other three trajectories were manually classified as others, but predicted as bursting by the LKCNN classifier. The accuracy percentages of each class are provided in the confusion matrix in Figure 8. The full classification results for all 126 time series, along with the curated data and the analysis scripts, are available in our GitHub repository. Note that the accuracy found in this section does not reflect the true performance of the LKCNN classifier, as the test dataset is highly curated. While the validation dataset may be subject to human errors, the true advantage of the LKCNN lies in its ability to operate beyond visually distinguishable data, providing fast and reliable classification. The results in the section provide an essential proof of concept, demonstrating not only how the model can be integrated into a data analysis pipeline but also how well it performs when confronted with the variability and noise inherent in experimental observations.

**Figure 7:**
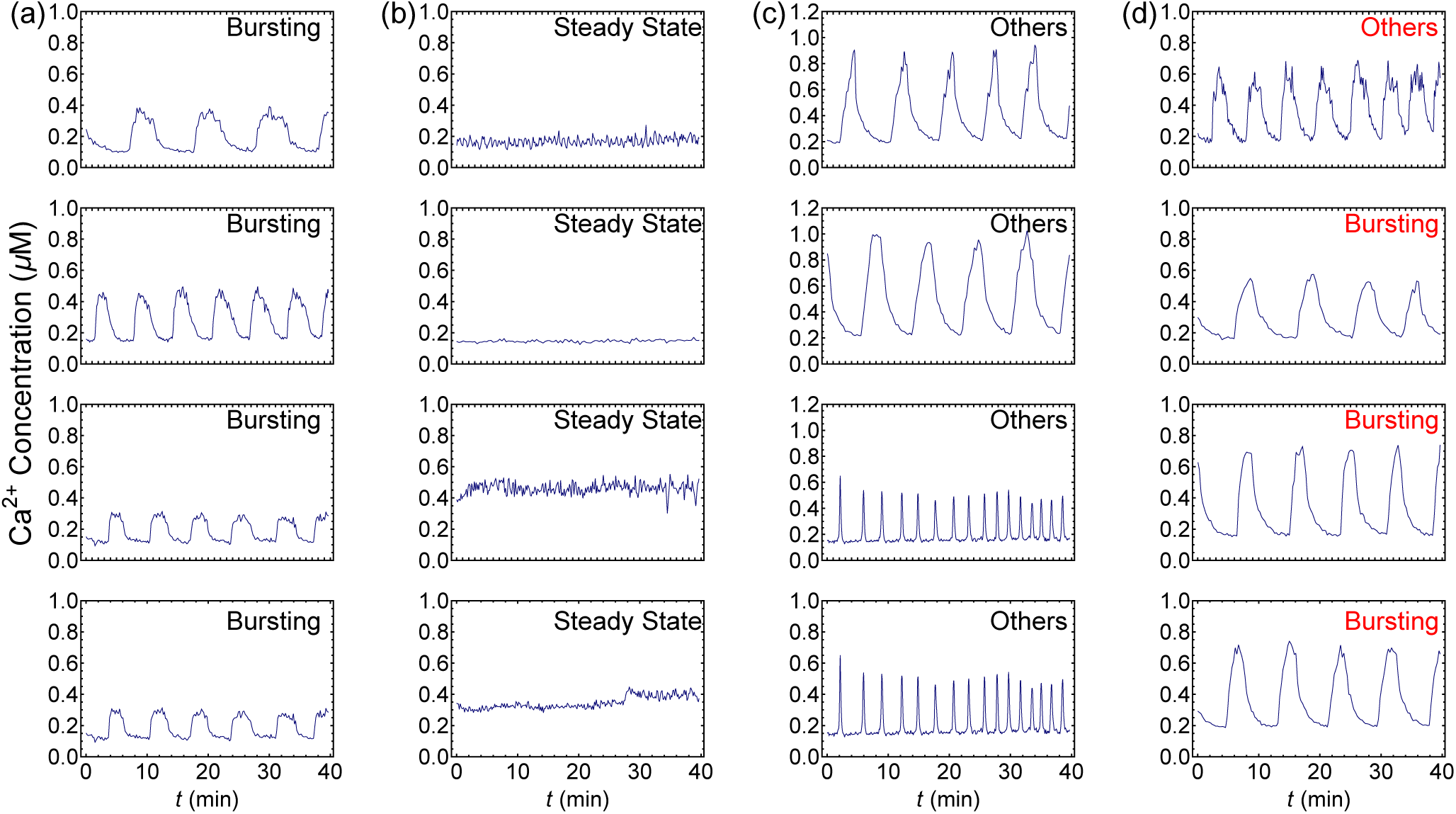
A sample of experimental trajectories with the LKCNN predicted class. Correctly classified (a) Bursting, (b) Steady States, (c) Oscillatory states. (d) Misclassified states where the first state was manually labelled as Bursting, while the other three were labelled as Others.

**Figure 8:**
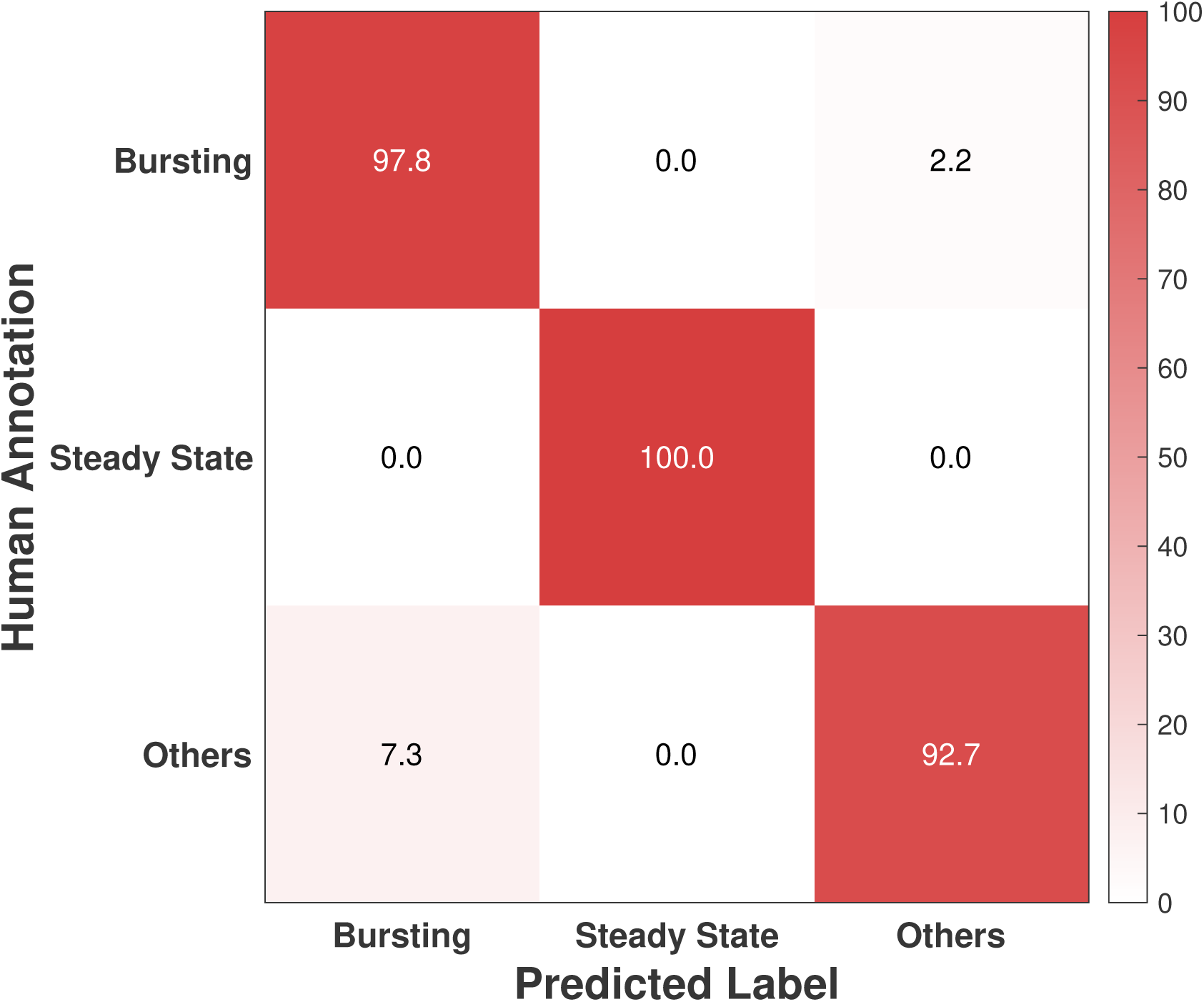
Confusion matrix showing the classification performance of our LKCNN model for the experimental dataset. Diagonal entries denote the percentage of correctly predicted dynamical states based on human annotation. Labels indicate three representative classes: Bursting, Steady State, and Others.

## DISCUSSION

Biological rhythms are central to life^47^. Some examples of biological oscillations include circadian rhythms, ovarian cycles, hormonal rhythms, mitotic cycles, and calcium oscillations^78^. Long-standing questions as to how we can decode the fluctuations in these physiological rhythms for meaningful information (for instance, patterns of the rhythms or anomalies in rhythmic time periods) that can help in a deeper understanding and better diagnosis of human disease remain active areas of research. Deep learning techniques combined with biomedical research can transform our understanding of the rhythms of life. Our present work is motivated by this research direction. This paper presents a proof-of-concept in applying a deep learning technique known as LKCNN to learn diverse patterns in the temporal dynamics of intracellular Ca^2+^ concentration that hold important physiological roles in cell biology.

In the present study, we establish the capability of the LKCNN framework to classify a broad range of intracellular Ca^2+^ dynamical regimes with high reliability. By extending the task from binary classification to an eight-class problem, the model captures the diversity of dynamical behaviors that underlie calcium signaling. Systematic evaluation of kernel size reveals that intermediate temporal receptive fields are optimal for balancing sensitivity to short-lived fluctuations with the ability to recognize longer-range temporal dependencies. These findings emphasize the architectural flexibility of LKCNNs in extracting features from complex biological time series.

Our analysis also highlights the impact of intrinsic stochasticity on classification accuracy. Whereas steady state, bursting, and simple oscillatory regimes were robustly distinguished, multiple-periodicity and chaotic dynamics were more difficult to resolve under elevated noise. This difficulty reflects a genuine overlap in their temporal signatures, making a clear separation a fundamental challenge. Indeed, a previous study using statistical measures confirmed this challenge, demonstrating that the distinction between these states diminishes under large intrinsic fluctuations^33^. While such measures are powerful for characterizing the complexity of known states, they often lack the clear decision boundaries required for robust classification. In contrast, our deep-learning approach is specifically designed for this classification task. Remarkably, it achieves an overall accuracy of over 90% even when the testing data set contained trajectories with noise levels two orders larger than its realistic value. This demonstrates that for the practical goal of automating the labeling of complex cellular dynamics, the feature-learning capabilities of the LKCNN framework offer a significant and robust advancement over traditional statistical methods.

Importantly, the application of LKCNN to experimental Ca^2+^ concentration traces demonstrated strong agreement with expert annotations, confirming its generalizability beyond synthetic datasets. This validation underscores the potential of LKCNNs as automated tools for high-throughput and unbiased classification of cellular dynamics. More broadly, the framework is well-positioned to extend to other oscillatory processes, including neuronal firing^79,80^, circadian rhythms^81,82^, and transcriptional dynamics^83^. Together, these results demonstrate the value of LKCNNs in bridging computational modeling and experimental biology, enabling scalable and systematic investigation of complex cellular behaviors.

### Limitations of the study

Although we have optimized the LKCNN classifier and achieved strong performance, our study still has two important limitations. The first, and most significant, is that we have assumed the system to remain stable within a single dynamical regime. In other words, our analysis has been conducted under the premise that the system’s underlying dynamics do not change during observation. In real experimental settings, however, this assumption often does not hold. For example, during an experiment, the external agonist, such as nutrient concentration, can be varied, which can introduce a shift or transition between different dynamical states over time. This means that a single time-series data generated experimentally can encompass multiple, distinct dynamical regimes. Such scenarios have not yet been addressed in the present work through our framework.

The second limitation of our study concerns the stability of the dynamical state in the presence of intrinsic fluctuations. Such fluctuations can drive the system toward instability, particularly when its parameters lie close to various phase boundaries. Systems exhibiting birhythmicity are particularly prone to this instability^24,84^. In these situations, the system may stochastically transition from one dynamical state to another. In our analysis, we deliberately excluded most of these cases by avoiding dynamical states that were near phase boundaries. However, it would be interesting to study the impact of such states on the performance of the classifier.

Another limitation of the study was the use of curated experimental data. In this study, the labels for the experimental data were manually annotated by researchers, relying on observation and subjective judgment. While this approach allowed for practical annotation during data collection, it also introduces potential bias and inconsistency, as human assessments can vary across individuals and contexts. An improved method would have been to derive the labels directly from physiological markers associated with each of the states of calcium concentration. Such an approach would provide objective, quantifiable evidence for distinguishing between conditions, thereby strengthening the validity of the dataset.

## METHODS

### Theoretical Nonlinear Model of Intracellular Calcium ion (Ca**^2+^**) Oscillations

We explain the mechanism of intracellular Ca^2+^ oscillations (panel (a) of Fig. 1) as follows. Consider a cell of system size represented by *V*. When an agonist attaches to the plasma membrane receptor, it triggers the synthesis of InsP_3_ (denoted by *Z*), another intracellular second messenger. InsP_3_ binds to receptors on the endoplasmic reticulum membrane, initiating Ca^2+^-release from the internal pool (denoted by *Y*) into the cytosol through the InsP_3_ receptor/Ca^2+^ channel (IP_3_R channel). This cytosolic Ca^2+^ (denoted by *X*) further activates its own release through the IP_3_R channel, a phenomenon known as Ca^2+^-induced Ca^2+^-release (CICR), indicating an autocatalytic process that generates intracellular Ca^2+^ oscillations. Stimulation of InsP_3_ 3-kinase activity is achieved through a Ca^2+^/calmodulin complex. Ca^2+^-activated InsP_3_ degradation occurs. In the figure, *β* represents the degree of cell stimulation by an agonist. Bold arrows encompass stimulus-induced Ca^2+^ influx, Ca^2+^ released from the pool, and Ca^2+^ exchange with the extracellular medium.

The intracellular Ca^2+^ oscillations can be described by a nonlinear theoretical model that couples free Ca^2+^ in the cytosol along with those of Ca^2+^ stored in the internal pool and InsP_3_. Suppose variables *X, Y*, and *Z* represent the populations of cytosolic Ca^2+^, Ca^2+^ stored, and InsP_3_, respectively. Then *x* = *X/V, y* = *Y/V,* and *z* = *Z/V* represent their respective concentrations. The time-evolution of these concentrations governs the complex dynamics of intracellular

Ca^2+ 24^ as,

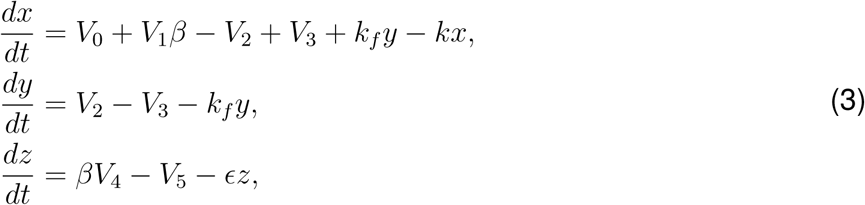

where

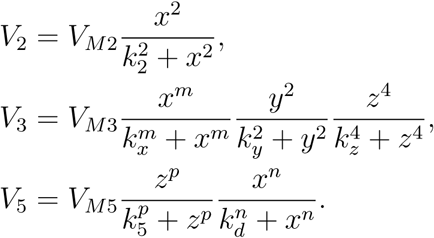

A detailed breakdown of the terms involved in Eq. (3) is as follows: *V*_0_ denotes the constant Ca^2+^ supply from the extracellular medium. *V*_1_ represents the maximum rate of stimulus-activated Ca^2+^ entry from the extracellular medium. The rate *V*_2_ (*V*_3_) corresponds to Ca^2+^ pumping from the cytosol into the internal pool (release of Ca^2+^ from the internal pool to the cytosol). *V_M_*_2_ and *V_M_*_3_ denote their maximum values. Parameters *k*_2_*, k_y_, k_x_*, and *k_z_* represent the threshold values for pumping, release, and activation of release by Ca^2+^ and InsP_3_, respectively. *V*_2_ is solely a function of the cytosolic Ca^2+^ concentration (*x*), whereas *V*_3_ depends on all the three concentrations *x, y*, and *z*. The rate constant *k_f_* measures the passive, linear leak of *y* into *x*, and *k* signifies the linear transport of cytosolic Ca^2+^ into the extracellular medium. *V*_4_ denotes the maximum rate of stimulus-induced InsP_3_ synthesis, and *V*_5_ represents the phosphorylation rate of InsP_3_ by the 3-kinase, an InsP_3_ metabolising enzyme. The decrease of InsP_3_ is driven by its hydrolysis by calcium-dependent 3-kinase. *k*_5_ denotes the half-saturation constant. Stimulation of InsP_3_ 3-kinase activity (through a Ca^2+^/calmodulin complex) is represented by a Hill-form term with *k_d_* as the threshold level of Ca^2+^. The term *ɛZ* accounts for the metabolism of InsP_3_ by 5-phosphatase, independent of Ca^2+^. Additionally, cooperative processes in Ca^2+^ release from internal stores into the cytosol and phosphorylation of InsP_3_ by 3-kinase are reflected in *V*_3_ and *V*_5_, incorporating Hill-coefficients *m*, *n* and *p*. A value of *p >* 1 indicates the presence of cooperativity in 3-kinase kinetics, while *p* = 1 indicates its absence^24^.

By adjusting the values of the rate constants and other model parameters in Eq. (3), concentrations *x*, *y*, and *z* exhibit various kinds of dynamical patterns^33^.

### Stochastic Modeling with Chemical Langevin Equation

Suppose *x*(*t*)*, y*(*t*)*, z*(*t*) represent the concentrations of cytosolic Ca^2+^, Ca^2+^ stored, and InsP_3_, respectively. Let the state vector of concentrations is *s* = *s*(*t*) = [*x*(*t*)*, y*(*t*)*, z*(*t*)]^T^. The evolution equation for the state, given previously in Eq. 3, can be re-written in a compact form as

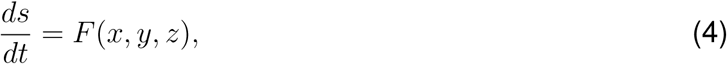

where

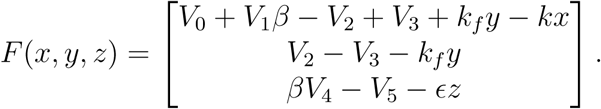

The system of ODEs in Eq. (4) can be reformulated as reaction channels^33^, highlighting the variation in populations *X, Y*, and *Z* as follows:

| Reaction channels | Propensity functions |
| --- | --- |
| $X \rightarrow X + 1$ | $VV_0$ |
| $X \rightarrow X + 1$ | $VV_1\beta$ |
| $X \rightarrow X - 1$ | $VV_2$ |
| $X \rightarrow X + 1$ | $VV_3$ |
| $X \rightarrow X + 1$ | $Vk_f y$ |
| $X \rightarrow X - 1$ | $Vkx$ |
| $Y \rightarrow Y + 1$ | $VV_2$ |
| $Y \rightarrow Y - 1$ | $VV_3$ |
| $Y \rightarrow Y - 1$ | $Vk_f y$ |
| $Z \rightarrow Z + 1$ | $V\beta V_4$ |
| $Z \rightarrow Z - 1$ | $VV_5$ |
| $Z \rightarrow Z - 1$ | $V\epsilon z$ |

In the second column, the propensity function for each reaction *R_j_*is calculated using Gillespie’s formalism^71,85^. The changes in the state *s* thus correspond to stochastic events involving the births and deaths of molecular species labeled as *X, Y*, and *Z*. These events inherently introduce randomness within the populations of these molecular species. Subsequently, we proceed to compute the CLE pertaining to the intracellular Ca^2+^ oscillation model (4) as follows:

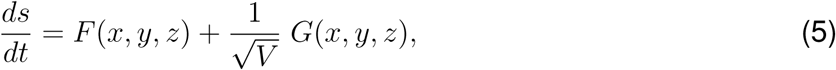

where *V* denotes the system size, and *G*(*x, y, z*) is:

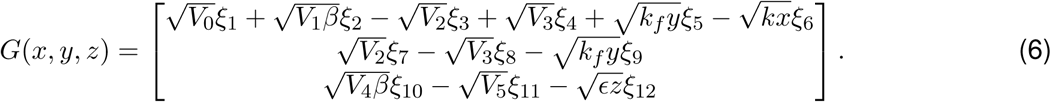

The stochastic differential equation (5) reduces to the deterministic equation (4), describing the mean-field behavior of the system, when *V→*∞

To simulate the intracellular Ca^2+^ dynamics, we numerically solve the CLE (1) using the *Euler-Maruyama* method, where the simulation is performed with a fixed time-step of 10*^−^*^6^ min. The volume parameter *V* is given in cubic micrometers (*µ*m^3^)^33^.

### Large Kernel Convolutional Neural Network (LKCNN)

For our Ca^2+^ dynamics classification task, the LKCNN architecture proceeds in such a way that the large kernel slides across the input time series to extract temporal features through successive convolutional layers and the extracted features are then flattened and processed through fully connected layers to produce the final classification output (as illustrated in Fig. 1(c)). This model architecture enables the network to capture both local temporal patterns and longrange dependencies.

The LKCNN classifier, we use a feed-forward architecture motivated by earlier approaches used for the classification of dynamical states^65,66^. The architectural details of the LKCNN utilized in our work are as follows:

- Convolutional Layer 1: 16 filters, kernel size *k*, stride 2 → Max Pooling (size 2)
- Convolutional Layer 2: 32 filters, kernel size *k*, stride 2 → Max Pooling (size 2)
- Dropout (rate 0.5) → Flatten
- Fully Connected Layers: 64 units → Dropout (rate 0.5) → 32 units → Output

The neural network uses ReLU activation functions for all hidden layers and is trained by minimizing the categorical cross-entropy loss function^86^ L given by:

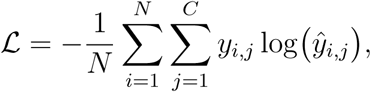

where *N* is the batch size, *C* is the number of classes, and *y_i,j_* is the ground-truth label. The term *y*^*_i,j_* represents the model’s predicted probability, calculated by applying the softmax function^86^ to the logits *z_i,j_* from the final output layer:

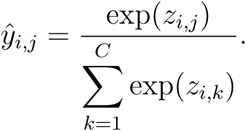

For optimization, we employ the Adam algorithm with a learning rate of *α*=10*^−^*^4^ and a batch size of *N* = 32. The training continues for at most 2, 000 epochs, incorporating an early stopping mechanism with a patience of 100 on the validation accuracy to prevent overfitting and select the best-performing model. To determine the optimal model configuration and ensure the robustness of our results, this entire procedure is systematically repeated for various kernel sizes, *k* 4, 5, 6*,…,* 100, and across 20 different random seeds. This rigorous exercise has identified an optimal kernel size of 28. The confusion matrices and final results presented in the main text are derived from the model trained using this optimal configuration.

### Parameters used for Synthetic Data Generation

**Table 1:** Parameter values used in the numerical simulation of the intracellular calcium oscillation model (1). Each region exhibits distinct dynamical behaviors: Region 1 (Bursting, Oscillatory, Steady State), Region 2 (Chaotic, Period-2, Period-3, Period-4, Steady State, Oscillatory), and Region 3 (Quasiperiodic, Steady State, Oscillatory). Within each region, 1000 synthetic trajectories were generated per dynamical pattern, each for training and testing the LKCNN classifier.

| Parameters | Region 1 | Region 2 | Region 3 |
| --- | --- | --- | --- |
| $\beta$ | [0.0, 0.8] | [0.65, 0.8] | [0.0, 0.8] |
| $\epsilon$ ($\text{min}^{-1}$) | [0.0, 10.0] | [10.0, 16.8] | [0.0, 2.5] |
| $V_0$ ($\mu\text{M min}^{-1}$) | 2 | 2 | 2 |
| $V_1$ ($\mu\text{M min}^{-1}$) | 2 | 2 | 2 |
| $V_{M2}$ ($\mu\text{M min}^{-1}$) | 6 | 6 | 6 |
| $k_2$ ($\mu\text{M}$) | 0.1 | 0.1 | 0.1 |
| $V_{M3}$ ($\mu\text{M min}^{-1}$) | 20 | 30 | 20 |
| $k_x$ ($\mu\text{M}$) | 0.3 | 0.6 | 0.5 |
| $k_y$ ($\mu\text{M}$) | 0.2 | 0.3 | 0.2 |
| $k_z$ ($\mu\text{M}$) | 0.1 | 0.4 | 0.2 |
| $V_{M5}$ ($\mu\text{M min}^{-1}$) | 30 | 50 | 30 |
| $k_s$ ($\mu\text{M}$) | 0.6 | 0.3194 | 0.3 |
| $k_d$ ($\mu\text{M}$) | 1 | 1 | 0.5 |
| $k_f$ ($\text{min}^{-1}$) | 10 | 1 | 1 |
| $k$ ($\text{min}^{-1}$) | 10 | 10 | 10 |
| $V_4$ ($\mu\text{M min}^{-1}$) | 3 | 2 | 5 |
| $m$ | 4 | 2 | 2 |
| $p$ | 1 | 1 | 2 |
| $n$ | 2 | 4 | 4 |

### Performance of Optimized Classifier

**Table 2:** Precision, recall, and F1 score for noiseless simulated trajectories of cytosolic Ca^2+^ concentration.

| Metric | Steady State | Bursting | Oscillatory | Period-2 | Period-3 | Period-4 | Chaotic | Quasiperiodic |
| --- | --- | --- | --- | --- | --- | --- | --- | --- |
| Precision | 1.0000 | 0.9999 | 0.9999 | 0.9756 | 0.9900 | 0.9930 | 0.9643 | 1.0000 |
| Recall | 1.0000 | 1.0000 | 0.9996 | 0.9957 | 0.9733 | 0.9691 | 0.9867 | 0.9999 |
| F1 score | 1.0000 | 0.9999 | 0.9998 | 0.9855 | 0.9815 | 0.9809 | 0.9753 | 0.9999 |

**Table 3:**
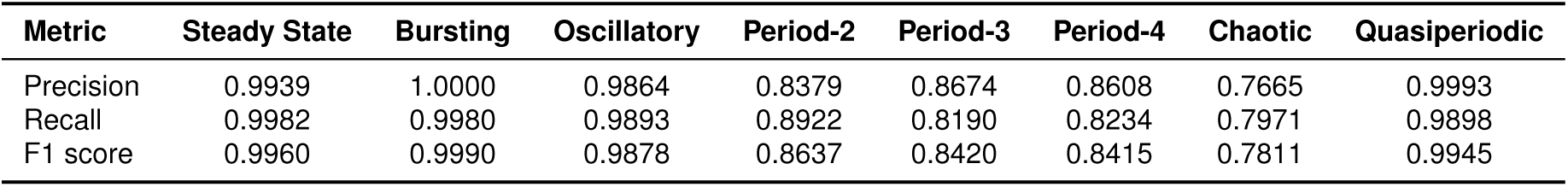
Precision, recall, and F1 score for noisy simulated trajectories of cytosolic Ca^2+^ concentration.

| Metric | Steady State | Bursting | Oscillatory | Period-2 | Period-3 | Period-4 | Chaotic | Quasiperiodic |
| --- | --- | --- | --- | --- | --- | --- | --- | --- |
| Precision | 0.9939 | 1.0000 | 0.9864 | 0.8379 | 0.8674 | 0.8608 | 0.7665 | 0.9993 |
| Recall | 0.9982 | 0.9980 | 0.9893 | 0.8922 | 0.8190 | 0.8234 | 0.7971 | 0.9898 |
| F1 score | 0.9960 | 0.9990 | 0.9878 | 0.8637 | 0.8420 | 0.8415 | 0.7811 | 0.9945 |

## RESOURCE AVAILABILITY

### Lead contact

Requests for further information and resources should be directed to and will be fulfilled by the lead contact, Jaesung Choi.

### Data and code availability

- All original code has been deposited at https://github.com/Jaesung-C/calcium_pattern_ lkcnn and is publicly available as of the date of publication.
- All original code or any additional information required to reanalyze the data reported in this paper is available from the lead contact upon request.

## ACKNOWLEDGMENTS

JC was supported by a KIAS Individual Grant (No. AP092902) via the Center for AI and Natural Sciences at the Korea Institute for Advanced Study (KIAS). SA was supported by a KIAS Individual Grant (No. PG096601) at the Korea Institute for Advanced Study (KIAS). ALC acknowledges research support from the YST and JRG programs at the APCTP through the Science and Technology Promotion Fund and Lottery Fund of the Korean Government (and local governments of Gyeongsangbuk-do Province and Pohang city). This work is supported by the Center for Advanced Computation at KIAS.

## AUTHOR CONTRIBUTIONS

Conceptualization, J.C, A.L.C. and S.A.; methodology, J.C., A.L.C. and S.A.; data curation, A.L.C. and S.A., investigation, J.C., and S.A.; Validation, J.C, A.L.C. and S.A.; writing-original draft, J.C. and A.L.C.; writing-–review & editing, S.A.; funding acquisition, J.C. and S.A.

